# Giant flagellins form thick flagellar filaments in two species of marine γ-proteobacteria

**DOI:** 10.1101/221366

**Authors:** Nicholas M. Thomson, Josie L. Ferreira, Teige R. Matthews-Palmer, Morgan Beeby, Mark J. Pallen

## Abstract

Flagella, the primary means of motility in bacteria, are helical filaments that function as microscopic propellers composed of thousands of copies of the protein flagellin. Here, we show that many bacteria encode “giant” flagellins, greater than a thousand amino acids in length, and that two species that encode giant flagellins, the marine γ- proteobacteria *Bermanella marisrubri* and *Oleibacter marinus*, produce monopolar flagellar filaments considerably thicker than filaments composed of shorter flagellin monomers. We confirm that the flagellum from *B. marisrubri* is built from its giant flagellin. Phylogenetic analysis reveals that the mechanism of evolution of giant flagellins has followed a stepwise process involving an internal domain duplication followed by insertion of an additional novel insert. This work illustrates how “the” bacterial flagellum should not be seen as a single, idealised structure, but as a continuum of evolved machines adapted to a range of niches.

## Introduction

Flagella are the organelles responsible for motility in diverse bacterial species. Flagella form helical filaments, several microns long, connected to a basal body that spans the cell envelope and functions as a rotary motor [1]. Some species have only a single flagellum, while others possess multiple filaments that form a coherent bundle for swimming [2]. Flagellar motors consist of over 20 different proteins that harness proton- or sodium-motive forces to generate torque [3]. The motor spins the flagellar filament to propel the cell.

The flagellar filament is composed of thousands of flagellin monomers whose architecture features a conserved polymerization core and a variable solvent-exposed region. Flagellins are secreted by the flagellar type III secretion system and travel through the hollow core of the growing filament to assemble into a helical array at the distal tip [4]. Flagellins from diverse species all possess conserved N- and C-terminal regions that discontinuously fold together to form the D0 and D1 structural domains [2,5]: D0 folds from the discontinuous extreme N- and C-terminal regions, while D1 folds from discontinuous adjacent N- and C-terminal regions. These domains mediate extensive inter-flagellin contacts, polymerizing to form the inner tubular section of the flagellar filament. The region between the discontinuous N-terminal and C-terminal segments of D1 forms the solvent-exposed surface of the filament, which, according to the structure of the canonical flagellin from *Salmonella enterica* serovar Typhimurium (*S.* Typhimurium), is composed of the D2 and D3 structural domains [5,6].

In contrast to the highly conserved N- and C-terminal flagellin domains, it has long been known that the central region of this protein is highly variable in both sequence and length [7]. Recent studies have also revealed considerable variation in flagellar motor structure together with alternative arrangements of variable surface-exposed flagellin domains that do not fit into the *S.* Typhimurium paradigm [2,8,9].

Most work on flagellins has focussed on proteins of a similar size to that of the phase 1 flagellin (FliC) of *S.* Typhimurium (495 amino acids; 51.6 kDa). However, nature provides multiple examples of ‘giant flagellins’ greater than a thousand amino acids in length, with large solvent-exposed hypervariable domains. Here, we sought to explore the phylogenetic and evolutionary context for these giant flagellins through bioinformatics analyses (also reviewed in our recent survey review [10]). We hypothesised that such giant flagellins might assemble into filaments thicker than the canonical flagellar filaments from *S.* Typhimurium and confirmed this experimentally by measuring filament thickness in two hydrocarbon-degrading marine γ-proteobacteria with predicted giant flagellins: *Bermanella marisrubri* Red65, originally isolated from the Red Sea [11] and *Oleibacter marinus* 2O1, originally isolated from Indonesian seawater [12]. We propose a mechanism for evolution of these giant flagellins based on phylogenetic analysis.

## Materials and methods

### Database searches and phylogenetic analysis of giant flagellins

A search of the September 2017 release of the UniProtKB database returned 4,678 protein sequences that harbour the PFAM domain pf00669 and are longer than *S.* Typhimurium FliC, including 92 longer than 1000 aa (S1 File). Although pf00669 represents the N-terminal helical structure common to all flagellins, it is also found in some representatives of the hook-filament junction protein FlgL. To extract only flagellins from the database, hidden Markov models (HMMs) for FliC and FlgL were constructed from established examples of each family (S2 and S3 Files). These HMMs were used in a voting scheme to identify 3536 true flagellins. To determine phylogenetic relationships, a multiple sequence alignment of the conserved N- and C-terminal flagellin sequences was built using FSA [13]. T-coffee was used to remove unconserved gaps [14]. A phylogeny was determined using RAxML [15], visualized with SeaView [16], and annotated with flagellin insert sizes using Inkscape 0.91 and Python scripting.

### Strains and growth conditions

Two strains potentially encoding giant flagellins were obtained as live cultures from the German Culture Collection (DSMZ; Table 1). They were grown in 5 mL of Marine Broth 2216 (BD, New Jersey, U.S.A.) supplemented after autoclaving with 1% filter-sterilised Tween 80, within loosely-capped 30 mL sterile polystyrene universal tubes at 28 °C and shaking at 200 rpm.

**Table 1.**
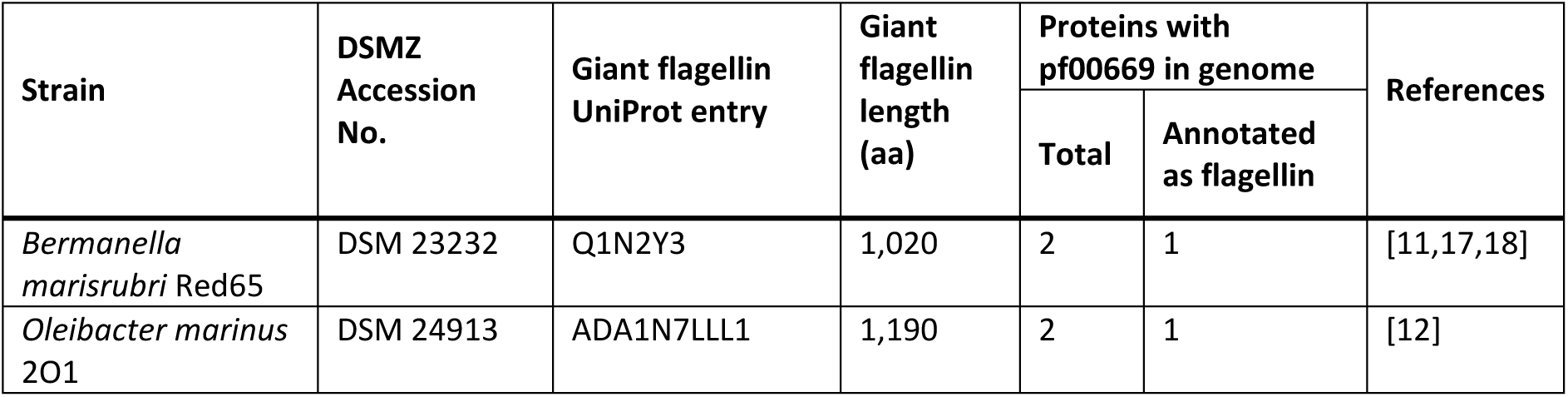
Bacterial strains used in this study.

### Light and transmission electron microscopy

Initial cultures were grown as described above for 72 h. A 50 µL portion of each culture was then transferred to 5 mL of the same culture medium and grown under the same conditions for 24 h. Motility was screened by placing a 5 µL drop of each culture on a microscope slide and examining with differential interference contrast through a 100× oil-immersion objective lens.

For transmission electron microscopy (TEM), 1 mL aliquots of the fresh cultures were washed twice by pelleting the cells at 1,500 × g for 3 min and gently re-suspending in 2-(N-morpholino)ethanesulfonic acid (MES) buffer. The final resuspension used 200 µL of MES buffer to achieve a higher cell density. 4 µL of the final cell suspension was deposited on glow discharged, carbon-coated TEM grids and the cells were allowed to settle onto the surface for 1 min. Cells were negatively stained by uranyl acetate and then imaged with a Tecnai T12 transmission electron microscope with TVIPS camera (FEI, Oregon, U.S.A.).

### Whole genome shotgun sequencing and analysis

*B. marisrubri* genomic DNA was purified from pelleted cells of a 5 mL culture using a FastDNA SPIN kit for Soil (MP Biomedicals, California, U.S.A.) according to the manufacturer’s instructions, except that the final elution used 100 µL of buffer DES. A sequencing library was prepared using a Nextera XT kit and sequenced on a NextSeq 500 in 2 × 150 bp, high-output mode (both Illumina, Cambridge U.K.).

Sequence reads were quality trimmed with Trimmomatic 0.38 [19] and assembled with SPAdes 3.11.1 [20]. Sequence similarity was assessed by calculating the average nucleotide identity between the resulting contigs and NCBI genome NZ_CH724113.1 at http://enve-omics.ce.gatech.edu/ani/index [21]. All reads were mapped to the expected flagellin gene sequence using the Geneious mapper and default parameters within Geneious 11.1.2. Mapped reads were separated and mapped again to the SPAdes contigs to locate the giant flagellin gene, and a pairwise alignment was created for the reference and test contig using MAFFT 7.388 [22].

### Protein identification by mass spectrometry

Flagella were isolated by acid depolymerisation according to an existing method [23], then separated by sodium dodecyl sulphate-polyacrylamide gel electrophoresis (SDS-PAGE) using 8% Novex Tris-Glycine gels (Fisher Scientific, Loughborough, U.K.). Following staining with Bio-Safe Coomassie Stain (Bio-Rad, Watford, U.K.), bands for analysis were cut out, washed, treated with trypsin, and extracted according to standard procedures [24]. The peptide solution resulting from the digest was mixed with α-cyano-4-hydroxycinnamic acid (Sigma-Aldrich, Gillingham, U.K.) as matrix and analysed on an Autoflex Speed MALDI-TOF/TOF mass spectrometer (Bruker Daltonics, Coventry, U.K.). The instrument was controlled by a flexControl method (version 3.4, Bruker Daltonics) optimised for peptide detection and calibrated using peptide standards (Bruker Daltonics). Data were processed in FlexAnalysis (Bruker Daltonics) and the peak lists were submitted for a database search using an in-house Mascot Server (v2.4.1; Matrixscience, London, UK). The search was performed against a custom database containing the sequence of interest (tr|Q1N2Y3|Q1N2Y3_9GAMM, Flagellin OS=Bermanella marisrubri GN=RED65_06563 PE=3 SV=1) in a background of approximately 200 random *E. coli* protein sequences using trypsin/P as enzyme with maximum 1 missed cleavage, 50 ppm mass tolerance, carbamidomethylation (C) as fixed and oxidation (M) as variable modification.

### Analysis of giant flagellin evolution

To analyse flagellin evolution, we focused on the *Oceanospiralles* and closely related species. We extracted sequences specific to the *Oceanospiralles* for further analysis (S1 File) and initially inspected sequences using Dotlet [25] to identify homologies and repeats. To identify the boundaries of the DE region, local sequence alignments were performed between pairs of tandem repeats from the same protein, and locally aligned parts of sequences extracted and used to build a multiple sequence alignment, as described above, of all DE regions that were found as tandem repeats (S4 File). An HMM was built from this multiple sequence alignment to identify remaining singly-occurring DE regions with appropriate boundaries. Matching sequences were extracted, pooled with the previously identified DE regions, and used to build a second HMM that was finally used to identify DE regions across the *Oceanospiralles* dataset (S1 Fig and S5 File). Another iteration did not identify additional regions, and no further iterations were performed.

To determine the scope of DE region occurrence, the DE region HMM was uploaded to the HMMSearch webserver version 2.23.0 at http://www.ebi.ac.uk/Tools/hmmer/search/hmmsearch and Reference Proteomes searched for significant hits [26].

## Results and discussion

### Giant flagellins are widespread in nature

Phylogenetic analysis (Fig 1) revealed that giant flagellins have evolved through convergent evolution within several different lineages scattered across a range of bacterial phyla and sub-phyla, including the α-, γ- and δ-/ ε- subdivisions of the Proteobacteria, Aquificae, Chrysiogenetes, Deferribacteres, Planctomycetes, Synergistetes and the Firmicutes, plus the candidate phylum Glassbacteria from the recently described candidate phyla radiation [27].

**Fig 1.**
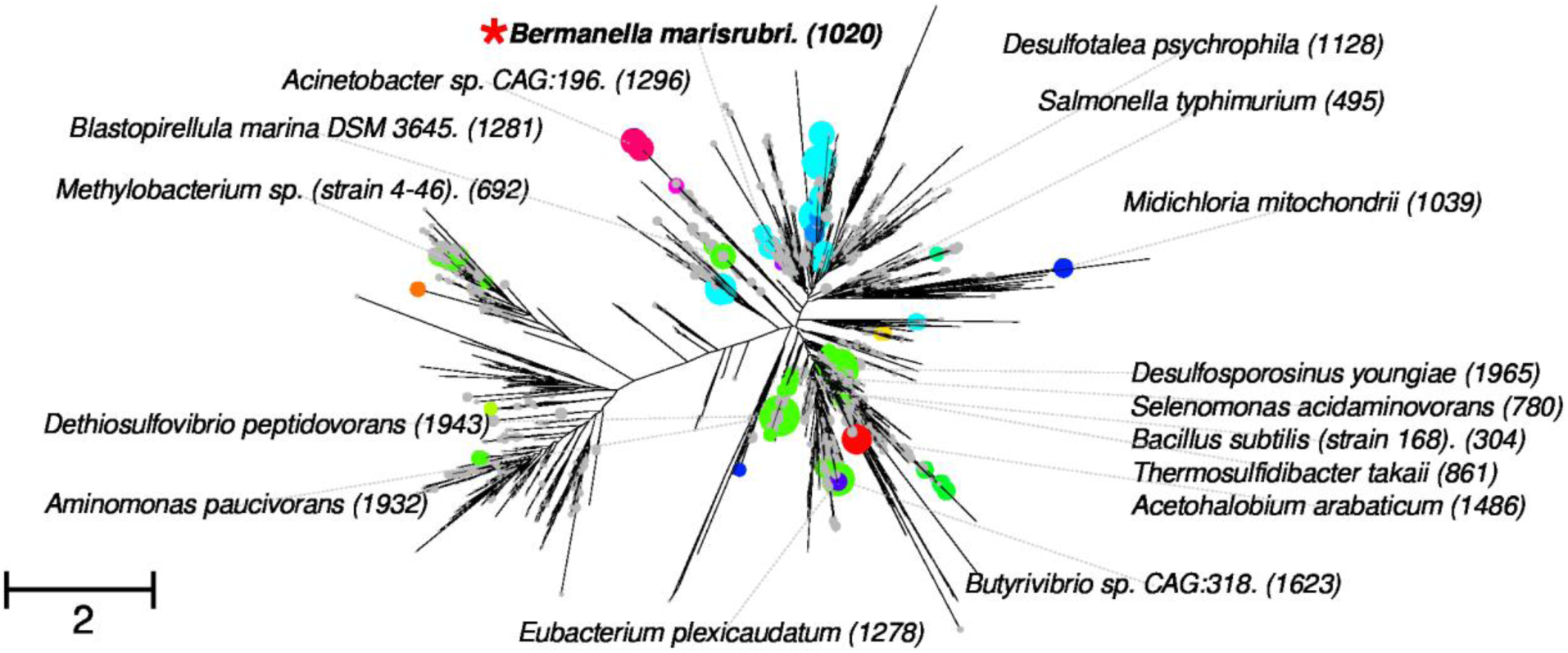
A phylogeny of giant flagellins. Phylogeny generated from conserved D0/D1 domain alignments; circles at ends of branches depict the size of the insertion, labels denote species with length of the full flagellin gene product in parentheses. Colours of circles represent discrete homology groups. The red star denotes the region in which *B. marisrubri* and *O. marinus* are found. Scale bar denotes average number of amino acid substitutions per site. Species are representative of diverse phyla, including α-proteobacteria (*Methylobacterium* sp.), γ-proteobacteria (*Acinetobacter* sp) and δ-/ ε- proteobacteria (*Desulfotalea psychrophila*), Aquificae (*Thermosulfidibacter takaii*), Planctomycetes (*Blastopirellula marina*), Synergistetes (*Aminomonas paucivorans*) and the Firmicutes (*Eubacterium plexicaudatum*).

### Giant flagellins produce thick flagella in the ***Oceanospiralles***

To determine whether giant flagellins produce thick flagella, two closely related γ- proteobacterial species with giant flagellins were selected for further investigation: *Bermanella marisrubri* Red65 and *Oleibacter marinus* 2O1. Both are representatives of the order *Oceanospiralles*, comprised of marine bacteria, many of which are able to consume petroleum hydrocarbons. The *B. marisrubri* (NCBI: NZ_CH724113.1) and *O. marinus* (NCBI: NZ_FTOH01000001.1) genomes each encode two proteins annotated as containing the pf00669 PFAM domain. However, in each case, the putative giant flagellin is the only protein predicted by our analyses to belong to the flagellin family, while the shorter proteins show greater similarity to FlgL than to flagellin.

Light microscopy revealed almost all *B. marisrubri* cells to be highly motile and to swim extremely quickly, while only a minority of *O. marinus* cells were motile (S1 and S2 Videos). Under the electron microscope, most *B. marisrubri* cells but far fewer *O. marinus* cells were flagellated, suggesting that differences in the numbers of motile cells reflected differential expression of the flagellar apparatus. *B. marisrubri* cells were very thin (< 0.5 µm), approximately 2 – 3 µm in length, with a spiral shape and a single, polar flagellum. Meanwhile, *O. marinus* cells were shorter (1 – 2 µm), curved or spiral in shape, also with a single, polar flagellum. In support of our hypothesis, we found that both *B. marisrubri* and *O. marinus* produced flagellar filaments with widths of at least 30 nm. This is approximately 25% thicker than *S.* Typhimurium phase I filaments (24 nm), the subunit of which contains 495 amino acids [5].

We confirmed the presence and sequence of the expected giant flagellin gene in *B. marisrubri* DSM 23232 by Illumina short-read whole genome shotgun sequencing (NCBI: SRP150336). Our assembled contigs had a 100% (S.D. 0.14%) two-way average nucleotide identity with the reference genome. The giant flagellin gene was located on contig 2 (333,297 bp). A pairwise alignment between this and the reference contig AAQH01000006 (146,124 bp) showed 99.9% identical bases, including the entire giant flagellin gene, with only an 84 bp intergenic sequence in the reference that was not in contig 2 (S2 Fig).

To confirm that in *B. marisrubri,* the giant flagellin is the only constituent of the flagellar filament, we isolated flagella from *B. marisrubri* cells by acid depolymerisation and detected a 120 kDa band by SDS-PAGE (S3 Fig). Tryptic digests of this protein band analysed by mass spectrometry produced peptides that matched with the expected protein (Q1N2Y3) in the UniProt database, with a protein score of 143 and E-value of 1.2 × 10^−12^ (S3 Fig and S6 File). The next-best match had a protein score of 18 with an E-value of 3.7, indicating a high degree of certainty over the identity of the expressed protein.

### Some giant flagellins contain repeat domains

Next, we sought to understand the evolutionary pathway to this giant flagellin family. Sequence analyses confirmed that the *B. marisrubri* flagellin features a large, non-homologous central insertion in place of the D2 and D3 domains found in *S.* Typhimurium. Dot plots applied to this central insert revealed an internal duplication within the relevant stretch of sequence from the giant flagellins seen in the *Oceanospiralles,* as exemplified by the flagellin from *B. marisrubri*. Analyses using Hidden Markov models of the repeat unit (which we have named the Domain-Extension or DE region to make lack of homology to D2 and D3 explicit) showed that this region was common in predicted flagellins from *Oceanospiralles* and related γ*-*proteobacteria. Large (>800 aa) and giant (>1,000 aa) flagellins from taxa closely related to *B. marisrubri* possess two tandem DE regions, while related *Oceanospiralles* species with intermediate-length flagellins feature only one DE region (Fig 2C). In addition to the tandem DE regions, *Oceanospiralles* with particularly giant flagellins (*Oleibacter, Thalassolituus, Oleispira*, and *Gynuella* species) feature an additional insert (which we call the DE-eXtension or DX insert), between the two DE regions. The phylogeny of post-duplication DE regions were congruent with the flagellin tree, suggesting a direct domain duplication followed by vertical transmission, as opposed to horizontal transfer of the insert.

**Fig 2.**
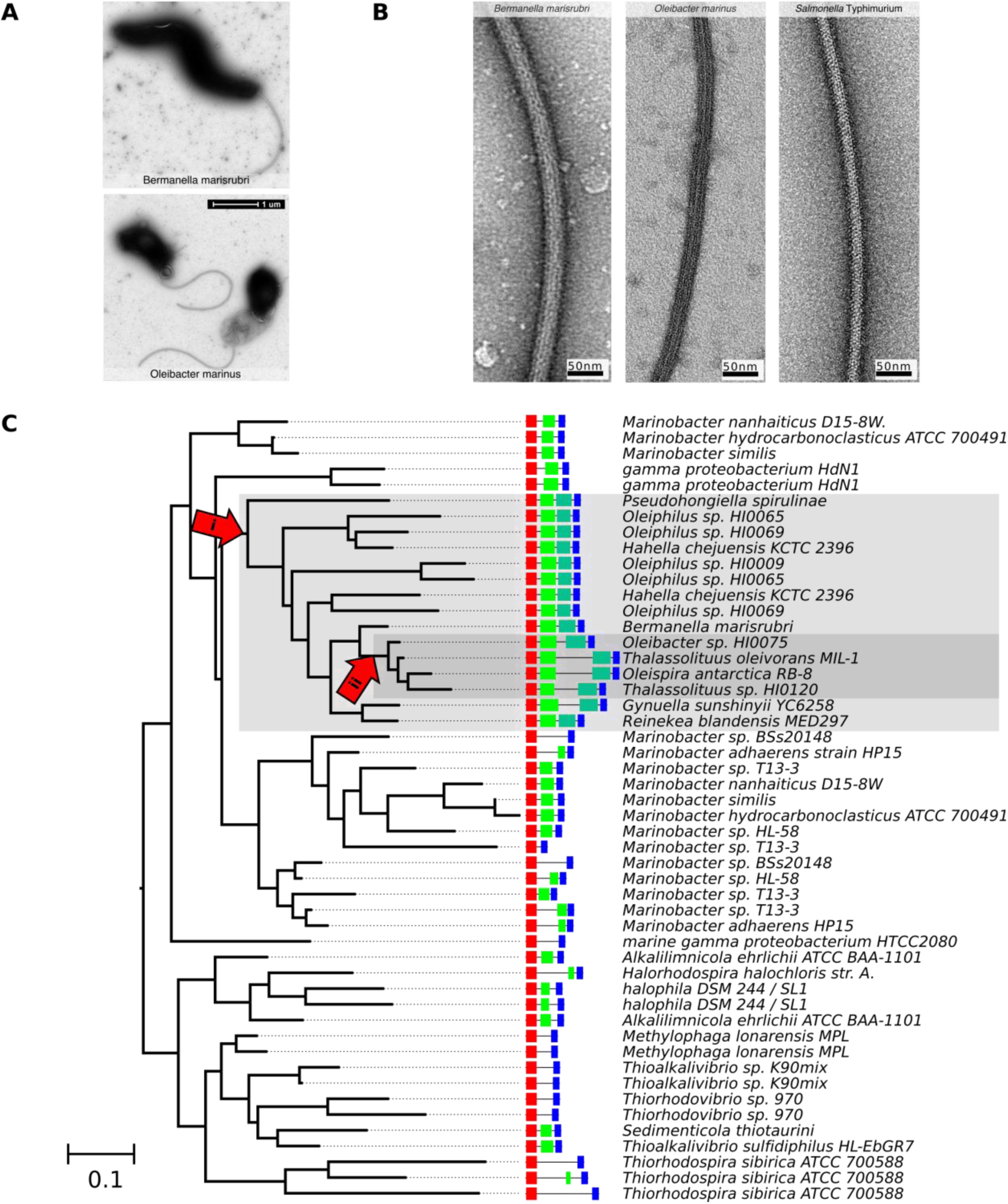
The *Oceanospiralles* giant flagellin forms a thick filament and evolved via an internal duplication. (A) TEM images of *B. marisrubri* and *O. marinus*. The images are typical of the cell morphology and flagellation pattern that we observed for each strain. (B) Negatively stained TEM images of representative flagellar filaments from *B. marisrubri* and *O. marinus*. A filament (24 nm diameter) from *S.* Typhimurium is also shown for comparison. All images are to the same scale. (C) Schematic depicting the phylogeny and occurrence of DE regions in related organisms. The phylogeny was determined using only the N- and C-terminal sequences that make up the D0 and D1 domains. Domain Extension (DE) regions are depicted in green; the C-terminal duplication is depicted in teal green. Arrows and accompanying grey boxes indicate inferred points at which the DE region was duplicated (“i”) and at which an additional insert was added between DE_1_ and DE_2_ (“ii”). Scale bar denotes average number of amino acid substitutions per site.

To determine the wider pattern of occurrence of the DE region, we searched Reference Proteomes using our hidden Markov model and HMMSearch. Significant hits were made to diverse bacteria including *Pseudomonas, Legionella, Campylobacter* and *Helicobacter* species, but not *Salmonella*. No significant hits were observed for any proteins other than annotated flagellins (E-value < 0.1). While partial structures of the *Pseudomonas* [2] and *Legionella* [28] flagellins have been determined, the structure of the DE region itself has not been determined to high resolution. The medium-resolution structure of the Pseudomonas DE domain region, however, is suggestive of inter-monomer pairwise dimerization of DE domains along the length of each of the 11 protofilaments [2].

The origin of the additional DX insert between tandem DE regions was less clear. Extraction of this sequence and BLASTing against reference genomes revealed no significant hits except among the *Oceanospiralles* flagellins. This DX insert features a conserved ∼35 aa glycine-rich region without internal repeats, which due to lack of other detectable homologs has probably been generated *de novo* [29].

How are the tandemly duplicated DE inserts likely to fold? Flagellins form discontinuous “out-and-back” topologies, wherein the N-terminus is close to the axis of the filament, the amino acids at the sequence midpoint are furthest from the flagellum axis and most peripheral, while the C-terminus is again axial, adjacent to the N-terminus. How, then, a flagellin can fold with a tandemly duplicated domain is intriguing. Two possibilities present themselves: the two DE regions fold internally, forming two tandem continuous DE domains corresponding to continuous stretches of sequence (Fig 3A), or the two DE repeats discontinuously fold to each contribute to one half of the two DE structural domains, as is the case with the D0, D1, and D2 domains (Fig 3B) in *S.* Typhimurium. Dimerization of the DE region of *Pseudomonas aeruginosa* [2], suggests that a domain duplication would lead to dimerization of the tandem DE regions, in favour of the former scenario, depicted in Fig 3A. Future structural determination will shed more light on the folding mechanism and evolution of these domains.

**Fig 3.**
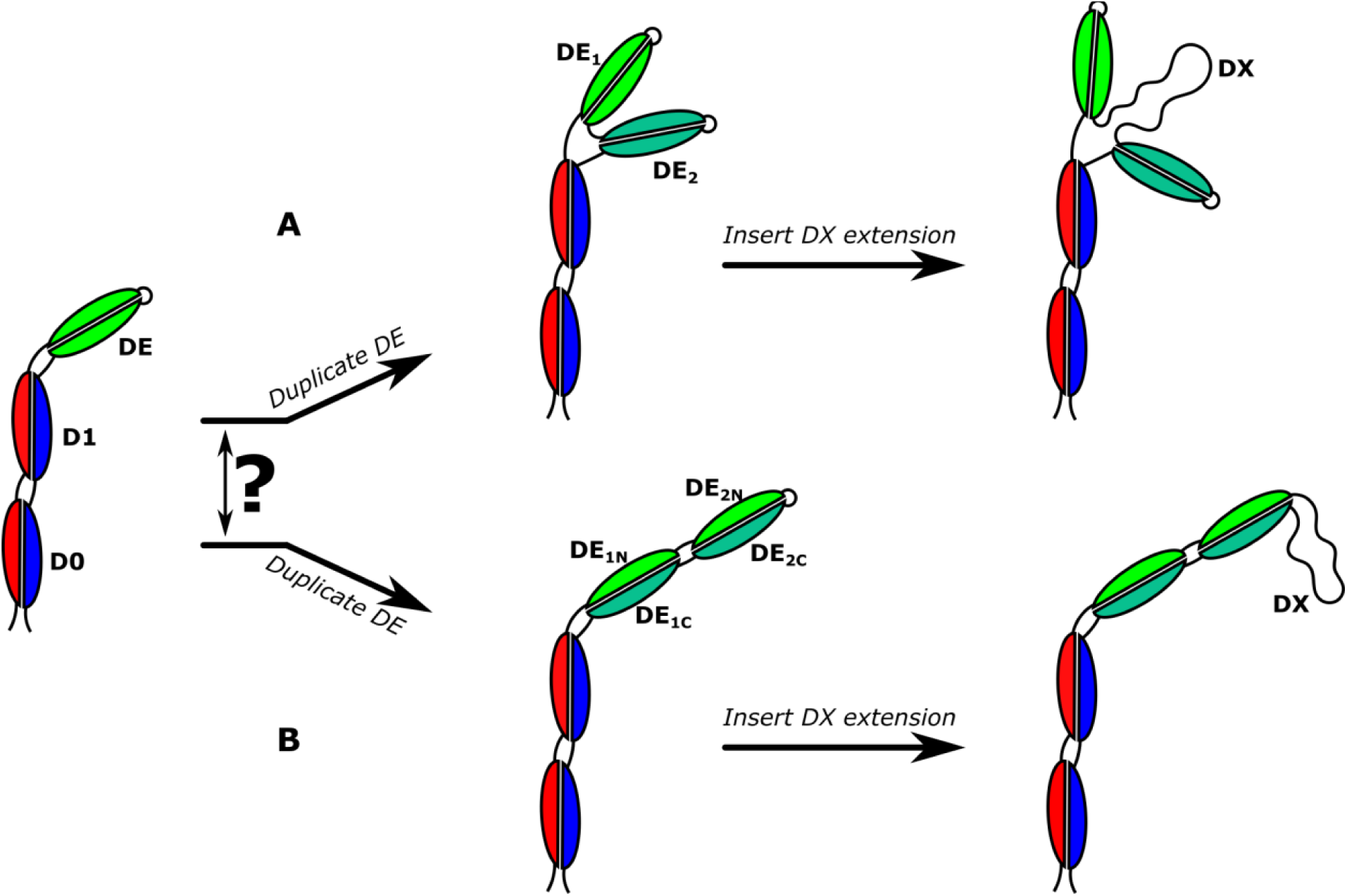
A model for evolution of the *Oceanospiralles* giant flagellin. Two possible scenarios for protein folding are conceivable: (A) the DE region duplicates and the two discrete DE sequences fold into discrete domains. Subsequent generation of the DX insert would make a more bulky flagellin. (B) Alternatively, the DE region duplication leads to two split domains, wherein the two physical DE regions are comprised of disparate parts of the two contiguous DE sequences. Subsequent insertion of the DX insert would make a longer protrusion at the surface.

### The functional advantages of giant flagellins and thick flagella remain unclear

Our phylogenetic analysis suggests an incremental, stepwise, and monotonic increase in flagellin size in the *B. marisrubri* and *O. marinus* lineage, indicating retention of each successively larger flagellin, and in turn suggesting consistent selective benefit from each size increase. We can currently only speculate as to the true driving force behind the evolution of giant flagellins. However, the parallel evolution in different— but not all—members of the *Oceanospiralles* indicates adaptation to a similar niche in each case. More broadly, the diversity of variable domains suggests that the protein has undergone similar processes of incremental adaptation to a myriad of different niches throughout the bacteria.

Multiple functions of flagellins also appears to be common: flagellins with active metallopeptidase enzyme domains have recently been discovered [30], and flagellins of enteric bacteria and plant pathogens also serve important functions in cell attachment and recognition or modulation of host immunity [1,31]. Therefore, it would appear that flagellins evolve to suit the specific combination of functional requirements demanded by their ecological niche.

The expanded central domain of the giant flagellins could confer a second function beyond motility, as described above. However, the proteins reported here do not possess sequence similarity to any enzyme domains. An alternative explanation is the increased thickness represents a mechanical adaptation to allow faster or more efficient swimming. In particular, a thicker flagellin might be less flexible due to steric hindrance, producing a longer, less helical filament. Motility at low viscosities might favour selection of such a less flexible filament as it would increase the axial thrust per rotation. Some species of bacteria, including *Vibrio parahaemoliticus* and *Bradyrhizobium japonicum* display a ‘division of labour’ by expressing single, thick, polar or subpolar flagella for swimming motility in liquid media and multiple, thinner, lateral flagella for swarming on surfaces or in viscous media [32–35]. However, *B. marisrubri* and *O. marinus* are not predicted to express any other flagellins and so probably rely solely on thick flagella for their motility.

The discovery of thick, polar flagellar filaments in two species of marine bacteria expands the known diversity of flagellar architecture, but also raises more questions. Are other components of the flagellar apparatus correspondingly large? Do thick flagella require alternative structural arrangements to those seen in the common model organisms? How common or widespread is this pattern of flagellation?

In conclusion, this work illustrates how “the” bacterial flagellum is not a single, ideal structure, but is instead a continuum of evolved machines adapted to a wide range of niches [36–38]. It will be interesting to learn how the structure and function of these marine flagellar systems are related to their sequences and how the structures are influenced by their environments.

## Acknowledgements

The authors would like to thank Dr Gerhard Saalbach (Proteomics Facility, John Innes Centre, U.K.) for his assistance with protein identification by mass spectrometry.

## Supporting information

**S1 File. Details of flagellins and strains used in the sequence analyses.** (Tab 1) Details of all flagellins with >495 amino acids (the length of *S.* Typhimurium FliC) in the September 2017 release of UniProtKB. (Tab 2) Selected *Oceanospiralles* flagellins and species used to identify sequence homologies and repeats.

**S2 File. Alignment of true flagellins used to build our hidden Markov model for distinguishing between flagellins and hook/hook-filament function proteins.** The file is a plain text file in ClustalX format that can be opened in any text editor for viewing.

**S3 File. Alignment of flagellar hook/hook-filament junction proteins (FlgE/FlgL) used to build our hidden Markov model for distinguishing between flagellins and hook/hook-filament function proteins.** The file is a plain text file in ClustalX format that can be opened in any text editor for viewing.

**S4 File. Multiple sequence alignment of all *Oceanospiralles* Domain Extension regions that were found as tandem repeats.** “ins” refers to the “insert” sequence, and “1half” and “2half” denotes either the first or second repeats. The file is a plain text file in ClustalX format that can be opened in any text editor for viewing.

**S1 Fig. Graphical logo representation of the Hidden Markov Model used to identify Domain Extension regions across the *Oceanospiralles.***

**S5 File. Original Hidden Markov Model used to identify Domain Extension regions across the *Oceanospiralles***. This file is in the standard output format from the HMMER suite of programs for Hidden Markov Model-based sequence alignment, available at www.hmmer.org.

**S1 Video. Swimming motility of *Bermanella marisrubri* Red65.** Cells were grown for 24 h in 5 mL of Marine Broth 2216 (BD, New Jersey, U.S.A.) supplemented after autoclaving with 1% filter-sterilised Tween 80, within loosely-capped 30 mL sterile polystyrene universal tubes at 28 °C and shaking at 200 rpm. A 5 µL sample was viewed using differential interference contrast and a 100× oil-immersion lens. The video playback speed has not been altered.

**S2 Video. Swimming motility of *Oleibacter marinus 2O1.*** Cells were grown for 24 h in 5 mL of Marine Broth 2216 (BD, New Jersey, U.S.A.) supplemented after autoclaving with 1% filter-sterilised Tween 80, within loosely-capped 30 mL sterile polystyrene universal tubes at 28 °C and shaking at 200 rpm. A 5 µL sample was viewed using differential interference contrast and a 100× oil-immersion lens. The video playback speed has not been altered.

**S2 Fig. Pairwise alignment of the flagellin-containing contigs in our genome assembly and the reference assembly.** (A) Overview of the whole alignment. (B) Detailed view of the alignment at the flagellin region (magenta box) showing 100% local identity. (C) Detailed view of the only sequence disagreement: an 84 bp intergenic region (olive box) present in the reference but not our assembly. The giant flagellin gene is represented by a magenta arrow. Other coding sequences are represented by yellow arrows.

**S3 Fig. Isolation and identification of the giant flagellin protein from *Bermanella marisrubri* Red65.** (A) SDS-PAGE gel of proteins isolated from the surface of *B. marisrubri* by acid depolymerisation, as described in materials and methods. The arrow indicates the 120 kDa band that was cut out and used for identification by trypsin digestion and mass spectrometry. (B) Summary of proteins identified by comparison of peptide fingerprints against a custom Mascot database. Protein score is −10 × Log(P), where P is the probability that the observed match is a random event. Protein scores outside of the green hatched area (>36) are significant (p<0.05).

**S6 File. Full Mascot search results for peptides generated by tryptic digest of the 120 kDa SDS-PAGE band.** Peptides were analysed by MALDI-TOF/TOF mass spectrometry. The top hits were to the expected *Bermanella marisrubri* giant flagellin.

## References

1. Rossez Y, Wolfson EB, Holmes A, Gally DL, Holden NJ. Bacterial flagella: twist and stick, or dodge across the kingdoms. PLoS Pathog. 2015;11: e1004483. doi:10.1371/journal.ppat.1004483

2. Wang F, Burrage AM, Postel S, Clark RE, Orlova A, Sundberg EJ, et al. A structural model of flagellar filament switching across multiple bacterial species. Nat Commun. 2017;8: 960. doi:10.1038/s41467-017-01075-5

3. Chen S, Beeby M, Murphy GE, Leadbetter JR, Hendrixson DR, Briegel A, et al. Structural diversity of bacterial flagellar motors. EMBO J. 2011;30: 2972–2981. doi:10.1038/emboj.2011.186

4. Paradis G, Chevance FF V., Liou W, Renault TT, Hughes KT, Rainville S, et al. Variability in bacterial flagella re-growth patterns after breakage. Sci Rep. Springer US; 2017;7: 1282. doi:10.1038/s41598-017-01302-5

5. Yonekura K, Maki-Yonekura S, Namba K. Complete atomic model of the bacterial flagellar filament by electron cryomicroscopy. Nature. 2003;424: 643–650. doi:10.1038/nature01830

6. Samatey FA, Imada K, Nagashima S, Vonderviszt F, Kumasaka T, Yamamoto M, et al. Structure of the bacterial flagellar protofilament and implications for a switch for supercoiling. Nature. 2001;410: 331–337. doi:10.1038/35066504

7. Beatson SA, Minamino T, Pallen MJ. Variation in bacterial flagellins: from sequence to structure. Trends Microbiol. 2006;14: 151–155. doi:10.1016/j.tim.2006.02.008

8. Beeby M, Ribardo DA, Brennan CA, Ruby EG, Jensen GJ, Hendrixson DR. Diverse high-torque bacterial flagellar motors assemble wider stator rings using a conserved protein scaffold. Proc Natl Acad Sci. 2016;113: E1917–E1926. doi:10.1073/pnas.1518952113

9. Beeby M. Motility in the epsilon-proteobacteria. Curr Opin Microbiol. Elsevier Ltd; 2015;28: 115–121. doi:10.1016/j.mib.2015.09.005

10. Thomson NM, Rossmann FM, Ferreira JL, Matthews-Palmer TR, Beeby M, Pallen MJ. Bacterial Flagellins: Does Size Matter? Trends Microbiol. Elsevier; 2017;0. doi:10.1016/j.tim.2017.11.010

11. Pinhassi J, Pujalte MJ, Pascual J, González JM, Lekunberri I, Pedrós-Alió C, et al. Bermanella marisrubri gen. nov., sp. nov., a genome-sequenced gammaproteobacterium from the Red Sea. Int J Syst Evol Microbiol. 2009;59: 373–377. doi:10.1099/ijs.0.002113-0

12. Teramoto M, Ohuchi M, Hatmanti A, Darmayati Y, Widyastuti Y, Harayama S, et al. Oleibacter marinus gen. nov., sp. nov., a bacterium that degrades petroleum aliphatic hydrocarbons in a tropical marine environment. Int J Syst Evol Microbiol. 2011;61: 375–380. doi:10.1099/ijs.0.018671-0

13. Bradley RK, Roberts A, Smoot M, Juvekar S, Do J, Dewey C, et al. Fast statistical alignment. PLoS Comput Biol. 2009;5. doi:10.1371/journal.pcbi.1000392

14. Notredame C, Higgins DG, Heringa J. T-coffee: A novel method for fast and accurate multiple sequence alignment. J Mol Biol. 2000;302: 205–217. doi:10.1006/jmbi.2000.4042

15. Stamatakis A. RAxML version 8: A tool for phylogenetic analysis and post-analysis of large phylogenies. Bioinformatics. 2014;30: 1312–1313. doi:10.1093/bioinformatics/btu033

16. Gouy M, Guindon S, Gascuel O. Sea view version 4: A multiplatform graphical user interface for sequence alignment and phylogenetic tree building. Mol Biol Evol. 2010;27: 221–224. doi:10.1093/molbev/msp259

17. Kube M, Chernikova TN, Al-Ramahi Y, Beloqui A, Lopez-Cortez N, Guazzaroni M-E, et al. Genome sequence and functional genomic analysis of the oil-degrading bacterium Oleispira antarctica. Nat Commun. 2013;4. doi:10.1038/ncomms3156

18. Rivers AR, Sharma S, Tringe SG, Martin J, Joye SB, Moran MA. Transcriptional response of bathypelagic marine bacterioplankton to the Deepwater Horizon oil spill. ISME J. Nature Publishing Group; 2013;7: 2315–2329. doi:10.1038/ismej.2013.129

19. Bolger AM, Lohse M, Usadel B. Trimmomatic: A flexible trimmer for Illumina sequence data. Bioinformatics. 2014;30: 2114–2120. doi:10.1093/bioinformatics/btu170

20. Bankevich A, Nurk S, Antipov D, Gurevich AA, Dvorkin M, Kulikov AS, et al. SPAdes: A New Genome Assembly Algorithm and Its Applications to Single-Cell Sequencing. J Comput Biol. 2012;19: 455–477. doi:10.1089/cmb.2012.0021

21. Goris J, Konstantinidis KT, Klappenbach JA, Coenye T, Vandamme P, Tiedje JM. DNA-DNA hybridization values and their relationship to whole-genome sequence similarities. Int J Syst Evol Microbiol. 2007;57: 81–91. doi:10.1099/ijs.0.64483-0

22. Katoh K, Standley DM. MAFFT multiple sequence alignment software version 7: Improvements in performance and usability. Mol Biol Evol. 2013;30: 772–780. doi:10.1093/molbev/mst010

23. Tahoun A, Jensen K, Corripio-Miyar Y, McAteer SP, Corbishley A, Mahajan A, et al. Functional analysis of bovine TLR5 and association with IgA responses of cattle following systemic immunisation with H7 flagella. Vet Res. 2015;46: 1–15. doi:10.1186/s13567-014-0135-2

24. Shevchenko A, Tomas H, Havli J, Olsen J V, Mann M. In-gel digestion for mass spectrometric characterization of proteins and proteomes. Nat Protoc. 2007;1: 2856–2860. doi:10.1038/nprot.2006.468

25. Junier T, Pagni M. Dotlet: Diagonal plots in a Web browser. Bioinformatics. 2000;16: 178–179. doi:10.1093/bioinformatics/16.2.178

26. Finn RD, Clements J, Arndt W, Miller BL, Wheeler TJ, Schreiber F, et al. HMMER web server: 2015 Update. Nucleic Acids Res. 2015;43: W30–W38. doi:10.1093/nar/gkv397

27. Brown CT, Hug LA, Thomas BC, Sharon I, Castelle CJ, Singh A, et al. Unusual biology across a group comprising more than 15% of domain Bacteria. Nature. 2015;523: 208–211. doi:10.1038/nature14486

28. Tenthorey JL, Haloupek N, López-Blanco JR, Grob P, Adamson E, Hartenian E, et al. The structural basis of flagellin detection by NAIP5: A strategy to limit pathogen immune evasion. Science (80-). 2017;358: 888–893. doi:10.1126/science.aao1140

29. McLysaght A, Hurst LD. Open questions in the study of de novo genes: What, how and why. Nat Rev Genet. Nature Publishing Group; 2016;17: 567–578. doi:10.1038/nrg.2016.78

30. Eckhard U, Bandukwala H, Mans MJ, Marino G, Holyoak T, Charles TC, et al. Discovery of a proteolytic flagellin family in diverse bacterial phyla that assembles enzymatically active flagella. Nat Commun. 2017;8:521. doi:10.1038/s41467-017-00599-0

31. Pel MJC, Pieterse CMJ. Microbial recognition and evasion of host immunity. J Exp Bot. 2013;64: 1237–1248. doi:10.1093/jxb/ers262

32. McCarter LL. Dual flagellar systems enable motility under different circumstances. J Mol Microbiol Biotechnol. 2004;7: 18–29.

33. Atsumi T, McCarter LL, Imae Y. Polar and lateral flagellar motors of marine Vibrio are driven by different ion-motive forces. Nature. 1992;355: 182–184. doi:doi:10.1038/355182a0

34. Kanbe M, Yagasaki J, Zehner S, Göttfert M, Aizawa SI. Characterization of two sets of subpolar flagella in Bradyrhizobium japonicum. J Bacteriol. 2007;189: 1083–1089. doi:10.1128/JB.01405-06

35. Quelas JI, Althabegoiti MJ, Jimenez-Sanchez C, Melgarejo AA, Marconi VI, Mongiardini EJ, et al. Swimming performance of Bradyrhizobium diazoefficiens is an emergent property of its two flagellar systems. Sci Rep. Nature Publishing Group; 2016;6: 23841. doi:10.1038/srep23841

36. Snyder LAS, Loman NJ, Fütterer K, Pallen MJ. Bacterial flagellar diversity and evolution: seek simplicity and distrust it? Trends Microbiol. 2009;17: 1–5. doi:10.1016/j.tim.2008.10.002

37. Pallen MJ, Matzke NJ. From The Origin of Species to the origin of bacterial flagella. Nat Rev Microbiol. 2006;4: 784–790. doi:10.1038/nrmicro1493

38. Aizawa S-I. The flagellar world: Electron microscopic images of bacterial flagella and related surface structures. 1st ed. Academic Press; 2013.

